# The Effects of Multi-Colour Light Filtering Glasses on Human Brain Wave Activity

**DOI:** 10.1101/2023.12.21.572905

**Authors:** Katherine Boere, Olave E. Krigolson

**Author notes:** **Correspondence:** Katherine Boere, PO Box (1700). STN CSC, University of Victoria Victoria, BC, Canada, V8W 2Y2.

## Abstract

The prevalence of electronic screens in modern society has significantly increased our exposure to high-energy blue and violet light wavelengths. Accumulating evidence links this exposure to adverse visual and cognitive effects and sleep disturbances. To mitigate these effects, the optical industry has introduced a variety of filtering glasses. However, the scientific validation of these glasses has often been based on subjective reports and a narrow range of objective measures, casting doubt on their true efficacy. In this study, we used electroencephalography (EEG) to record brain wave activity to evaluate the effects of glasses that filter multiple wavelengths (blue, violet, indigo, and green) on human brain function. Our results demonstrate that wearing these multi-colour light filtering glasses significantly reduces beta wave power (13-30 Hz) compared to control or no glasses. Prior research has associated a reduction in beta power with the calming of heightened mental states, such as anxiety. As such, our results suggest that wearing glasses such as the ones used in this study may also positively change mental states, for instance, by promoting relaxation. This investigation is innovative in applying neuroimaging techniques to confirm that light-filtering glasses can induce measurable changes in brain activity.

---

The influence of blue light on our health has emerged as a significant point of interest in scientific research, mainly due to the widespread emission of blue light wavelengths from light-emitting diodes (LEDs), compact fluorescent lamps, and electronic devices with displays (Munsamy et al., 2022). In moderation, blue light is vital for maintaining visual health and ensuring optimal hormonal and cognitive function (Gehring & Rosbash, 2003; Mainster, 2006; Münch et al., 2016). However, the benefits of blue light are offset by the adverse effects of prolonged exposure, which include symptoms such as fatigue, reduced cognitive performance, and disrupted sleep patterns (Ide et al., 2015; Munsamy et al., 2022; Rosenfield, 2011). In today’s digitally immersed era, where screens are an essential part of daily life, exposure to blue light has reached extraordinary levels (Souchet et al., 2022). This increased exposure raises crucial questions about its impact on our neural and physiological systems and whether modern interventions, like light-filtering glasses, can effectively mitigate potential adverse effects.

Light-filtering glasses, created to mitigate the effects of excessive blue light exposure, are engineered with a specialized dye coating tailored to selectively block, absorb, or attenuate potentially harmful light waves (Palavets & Rosenfield, 2019). Notably, these glasses primarily target blue light while allowing other beneficial wavelengths, such as violet, indigo, and green, to pass through (Vagge et al., 2021). The rise in digital device usage has sparked a growing consumer demand for protective eyewear, propelling it to become a rapidly evolving market segment in the optical industry. Although manufacturers advocate for their products, claiming they can counteract the negative effects of blue light overexposure, the efficacy of blue light filtering glasses is still debated and calls for a more thorough understanding (Lawrenson et al., 2017; Munsamy et al., 2022; Vagge et al., 2021).

For instance, research by Leung et al. (2017) reported no discernible difference in cognitive performance among participants wearing high blue-light filtering glasses, those with low blue-light filtering glasses, and those with clear control glasses after a two-hour computer task. Similarly, Palavets and Rosenfield (2019) discovered that although blue-light blocking glasses filtered out 99% of short-wavelength light, their effectiveness in reducing eye strain and fatigue was not superior to that of a neutral control filter. These findings are supported by several meta-analytical reviews, including those by Lawrenson et al. (2017), Munsamy et al. (2022), and Vagge et al. (2021), which highlight consistent inconsistencies and the absence of robust objective measures in existing research on blue light filtering glasses. This emphasizes the urgent need to investigate further the direct impacts of these glasses on brain function.

Given the notable inconsistencies and gaps in the literature regarding the effects of light-filtering glasses, a more detailed approach is essential to elucidate the underlying neural mechanisms involved. Electroencephalography (EEG) is a non-invasive neuroimaging tool that measures the brain’s electrical activity through electrodes on the scalp (Cohen, 2017). The resulting waveforms, reflective of rhythmic neural oscillations, are categorized into frequency bands—delta (1 to 3 Hz), theta (4 to 7 Hz), alpha (8 to 12 Hz), and beta (13 to 30 Hz)—each associated with different mental states and cognitive processes (Egner, 2004; Knott et al., 1996; Klimesch, 1999; Angelakis et al., 2007; Cavanagh & Frank, 2014; Knyazev, 2012).

In this context, our study employed EEG to assess the impact of multi-colour light-filtering glasses on brain wave activity, specifically the oscillatory patterns in the brain corresponding to theta, alpha, and beta rhythms. We compared the effects of wearing light-filtering glasses that filter violet, indigo, blue, and green light to those of clear control glasses and no glasses on EEG oscillations. Our primary hypothesis posits that EEG oscillations will change when participants wear light-filtering glasses compared to the control conditions. Supported by limited literature suggesting that light-filtering glasses may reduce fatigue during exposure (Ide et al., 2015; Lin et al., 2017), we anticipated that participants would exhibit decreased frontal alpha oscillations while wearing the light-filtering glasses, along with potential changes in theta and/or beta oscillations. Understanding the effects of light-filtering glasses on brain activity carries significant implications for individual well-being and public health, potentially informing guidelines for the responsible use of electronic devices and energy-efficient lighting. Through this study, we aim to contribute to the expanding body of knowledge on the impact of blue light on human health and to promote the development of effective strategies to mitigate its possible adverse effects.

## Methods

### Participants

Forty participants (mean age 43 years old [age range 24-65], 25 female) participated in the study. Participants were recruited between June 1, 2022, and September 1, 2022. Eighteen participants required corrective glasses. In addition, each participant completed the Perceived Stress Test (PSS-10) before commencing the experiment (Cohen, 1994). Five participants were removed from the analysis due to excessive noise in their EEG data (actual n = 45). Participants all provided written and informed consent approved by the Human Research Ethics Board at the University of Victoria. Participants were asked not to consume caffeine or exercise the day of the experiment and not to consume alcohol within 24 hours of the experiment. All participants received a pair of TrueDark® Twilight filtering glasses for their participation in the study.

### Procedure

Participants were seated in a sound-dampened room with 60-watt fluorescent lighting. TrueDark® Twilight glasses were used in this experiment (TrueDark®; Washington, USA). These glasses are uniformly tinted and block 99.27% of violet, indigo, blue and green light, ranging between 380nm – 570nm (for a full lens transmittance report, see the Supplementary Information). Every participant’s session included four distinct five-minute EEG recording blocks, leading to a total data collection period of 20 minutes. Each recording block was sequentially paused, categorized, and saved under a unique identifier, ensuring participants’ anonymity. The procedure commenced with participants seated comfortably in a chair, where they were asked to relax as much as possible. Participants were directed to gently fix their gaze on a Post-it note adhered to a blank wall in front of them. Participants who needed prescription eyeglasses were instructed to keep them on throughout the experiment. The initial recording condition for all participants entailed capturing EEG data while their eyes were open in a resting state. Following this, participants proceeded to the second and third conditions, which involved wearing light-filtering or clear control glasses. The sequence of these conditions alternated between participants. For instance, the second condition for Participant 1 entailed wearing light-filtering glasses, followed by clear control glasses in the third condition.

Conversely, Participant 2 wore the clear control glasses in the second condition, followed by the light-filtering glasses in the third condition. There was a five-minute adaptation period between each lensed condition to mitigate the previous condition’s effects and prepare the eyes for the following testing condition (Rahman et al., 2014). Each session concluded with the final condition – an EEG recording with the participant’s eyes closed.

### Data Acquisition

EEG data were acquired using a CGX Quick-20 Dry EEG headset (Cognionics; CA, USA). The headset has 20 active dry sensors (plus two references and one ground) arranged according to the International 10–20 System. Each sensor is paired with an active amplifier and shield. The ground electrode was positioned on the forehead centred between Fp1 and Fp2. Reference electrodes (A1, A2) were attached to the left and right earlobes. Electrode impedances were kept below 2,500 kΩ to ensure optimal data quality. Data were recorded through a built-in amplifier and transmitted wirelessly via Bluetooth to the acquisition computer, where they were stored for offline analyses. Data were then digitized at 1 kHz using the Cognionics Data Acquisition 2.0 Software (http://cognionics.com/wiki/pmwiki.php/Main/DataAcquisitionSoftware).

### Data Processing

Processing and analysis were conducted using custom code MATLAB scripts that used EEGLAB (version 9.10.0.1739362 (R2022b)) environment (Delorme & Makeig, 2004) running on MATLAB 2022a (MathWorks Inc., Natick, USA) on Windows 10. All analysis code can be found at https://github.com/Neuro-Tools. First, each data set was filtered using a dual-pass Butterworth filter with a passband of 0.1–30 Hz (order two roll-off) and a notch filter of 60 Hz. Next, data were divided into temporal epochs of 1000ms segments with 500ms overlaps and run through artifact rejection, where trials with an absolute difference of 150 μV were removed. Data were then transformed using Fast Fourier Transform, the standard MATLAB function similar to Cohen (Cohen, 2014). Fast Fourier Transform results were then averaged over all epochs, and power was computed for each frequency band of interest. Specifically, we computed the average power for the theta (4 to 7 Hz), alpha (8 to 12 Hz) and beta (13 to 30 Hz) bands (i.e., the bands of interest) (μV^2^) at electrode Fz (the frontal electrode). As we were primarily interested in the cognitive states of attention, focus, and relaxation for this study, we excluded the delta band (1-3 Hz) from our analysis.

### Data Analysis

To investigate the difference between conditions, we conducted a repeated measure analysis of variance (ANOVA) for each averaged frequency band theta (4 to 7 Hz), alpha (8 to 12 Hz) and beta (13 to 30 Hz) across conditions. This analysis step was followed by a pairwise comparison using the Holm correction to investigate the difference between each condition, verifying the effects of light-filtering glasses on brain activity. Notably, all error bars on the figure represent 95% within-subject confidence intervals. All statistical analyses were conducted in R (Version 3.5.3; R Core Team, 2019).

## Results

The repeated measures ANOVA identified significant differences in beta band power across the conditions (clear control glasses, no glasses, and light-filtering glasses) (F(2,78) = 13.68, *p* < .0001), with sphericity assumed and confirmed. Pairwise comparisons revealed significant differences in beta power between the control condition (*M* = 0.75, *SD* = 0.11) and the light-filtering glasses condition (*M* = 0.65, *SD* = 0.095), *t*(78) = 4.234, *p* < .0001 (see Figure 1), and between the no glasses condition (*M* = 0.77, *SD* = 0.12) and the light-filtering glasses condition (*M* = 0.65, *SD* = 0.095), t(78) = 4.958, *p* < .0001 (see Figure 1). No significant differences were found in beta power between the clear glasses and no glasses conditions (p > .05). Likewise, no significant differences were observed in theta (F(2,78) = 1.525, *p* > .05) or alpha (F(2,78) = 0.728, *p* > .05) band power among the conditions. All statistical assumptions were tested and met.

**Figure 1.**
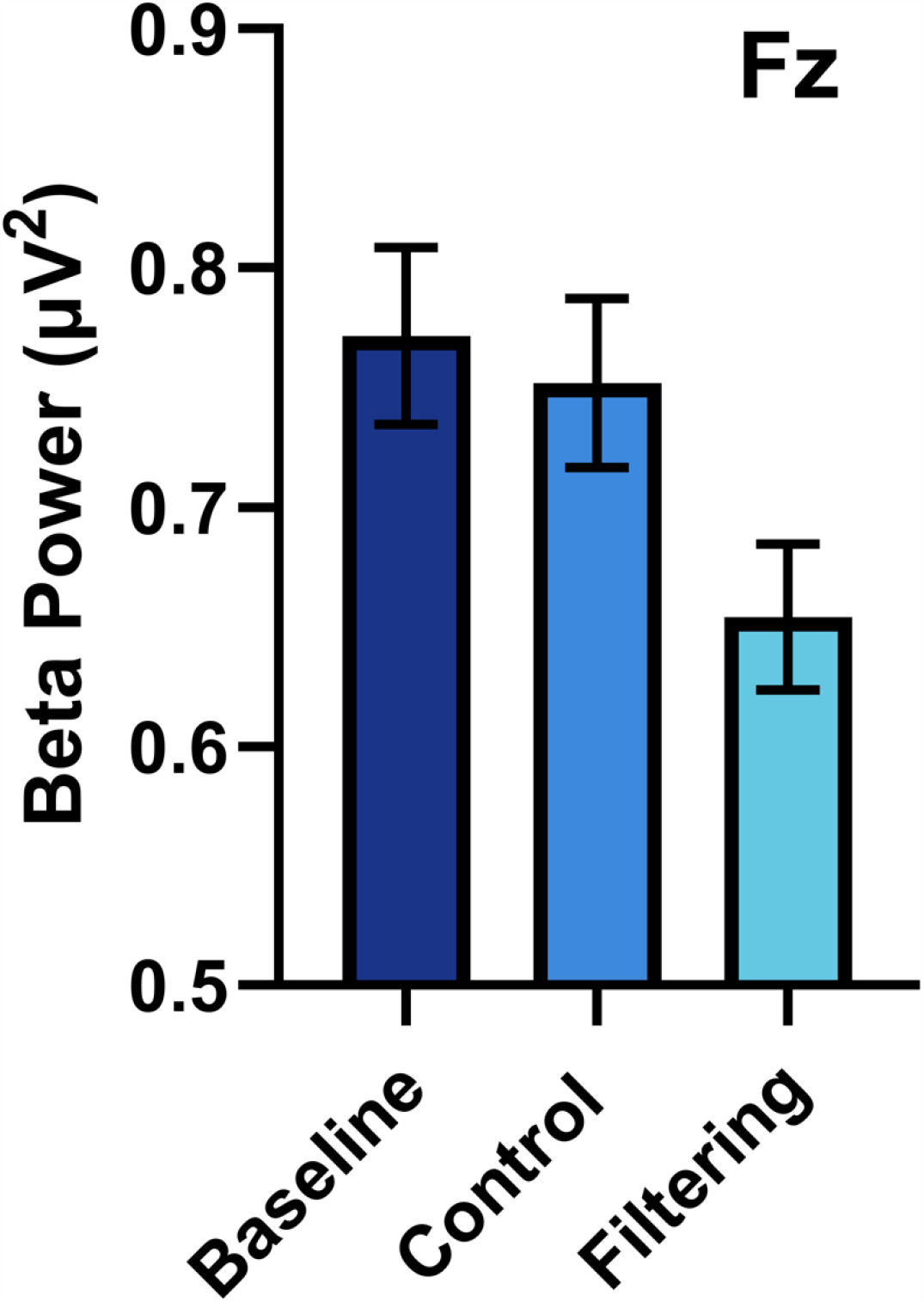
Mean frontal (Fz) beta power (13-30Hz) for baseline (no glasses), control (clear glasses) and filtering glasses. All error bars represent 95% confidence intervals.s

## Discussion

Here, we have demonstrated that wearing light-filtering glasses produces changes in the human EEG signal. Specifically, our results show that TrueDark® filtering glasses reduce frontal beta activity compared to clear control glasses or when wearing no glasses. This finding aligns with our primary hypothesis. However, we did not find evidence to support our secondary hypothesis, as no difference in frontal alpha power was recorded between conditions. Frontal alpha power has been previously tied to inner focus and concentration (Angelakis et al., 2007). Therefore, we propose this discrepancy is because participants’ EEG activity was recorded during the resting state and lacked a specific task to focus on apart from the directions to gaze softly at the fixation cross.

The key finding from our study is the decrease in frontal beta oscillations found while wearing light-filtering glasses. As mentioned, EEG band oscillations are commonly associated with numerous psychological states; however, *decreased* beta-band activity has been less readily examined (Engel & Fries, 2010). With that said a limited number of studies have suggested that decreased frontal beta oscillations are linked to increased relaxation (Diego et al., 2004; Ernst Niedermeyer & Silva, 2005; Hammond, 2005; Jacobs et al., 1996; Schoneveld, 2016). For example, Diego et al. (2004) utilized EEG to assess how therapeutic massage affects acute anxiety. Here, the researchers reported a positive association between reduced frontal beta activity and relaxation (Diego et al., 2004). In addition, Schoneveld and colleagues (2016) found that decreased frontal beta activity was associated with a decrease in symptoms of anxiety reported on the DSM-IV (a commonly used diagnostic tool for depression and anxiety disorders, see Bell, 1994) (Schoneveld, 2016). Together, these findings may suggest that the decrease in frontal beta power observed here when participants wore light-filtering glasses reflects a reduction in highly aroused or anxious states, thereby increasing relaxation (Ernst Niedermeyer & Silva, 2005).

Notably, there are constraints to the conclusions we can draw from our findings. We investigated how multi-colour light filtering glasses worn during resting state may impact brain activity. However, the found effects were not isolated by an experimental task, and we could not discern their influences elicited independently by the light-filtering glasses. For example, as mentioned above, decreased frontal beta activity has been linked to relaxation; however, other studies connect reduced beta activity to symptoms of ADHD (Arns et al., 2012). Thus, it could be that the light-filtering glasses caused the observed change in beta activity or that another factor caused it, such as decreased attention span as exhibited in ADHD, or it could be that these factors caused it interactively. Consequently, we cannot determine the exact mechanism of the observed shift in beta oscillations. We recommend that future research utilize tasks that can systematically control possible means to discern the light-filtering glasses’ specific contribution to changes in brain wave activity.

In sum, this study is the first to assess the effect of multi-colour light-filtering glasses on brain wave activity. Our results showed a more significant decrease in beta power while participants wore light-filtering glasses than clear control glasses during a five-minute resting state EEG recording. We have proposed that this indicates a reduction in highly activated mental states, such as anxiety, resulting in increased mental relaxation. Importantly, our findings provide objective support for the efficacy of light-filtering glasses, specifically ones that filter violet, indigo, blue and green light. This research then sets the groundwork for future neuroimaging tools to examine the brain changes caused by light-filtering glasses and the specific mechanisms behind these changes.

## Author Contributions

KB designed the study, collected the data, completed the analysis and wrote the manuscript. OK oversaw the project and contributed to the manuscript. All authors approved the submitted version.

## Funding

This research was supported by the NSERC Discovery Grant (RGPIN 2016-0943).

## Data Availability Statement

All processing scripts can be found at https://github.com/Neuro-Tools. In addition, the data supporting this study’s findings are available from https://osf.io/nj56y/.

## Notes

### Competing Interest Statement

The authors have declared no competing interest.

